# The dengue virus non-structural protein 1 (NS1) interacts with the putative epigenetic regulator DIDO1 to promote flavivirus replication

**DOI:** 10.1101/2021.09.01.458517

**Authors:** Gerson I. Caraballo, Romel Rosales, Mercedes Viettri, Siyuan Ding, Harry B. Greenberg, Juan E. Ludert

## Abstract

Dengue virus (DENV) NS1 is a multifunctional protein essential for viral replication. To gain insights into NS1 functions in mosquito cells, the protein interactome of DENV NS1 in C6/36 cells was investigated using a proximity biotinylation system and mass spectrometry. Approximately 14% of the 817 identified proteins coincide with interactomes obtained in vertebrate cells, including ontology groups of the oligosaccharide transferase complex, the chaperonin containing TCP-1, and nuclear import and export, vesicle localization and ribosomal proteins. Notably, other protein pathways such as epigenetic regulation and RNA silencing, not previously reported in vertebrate cells, were also found in the NS1 interactome in mosquito cells. Due to the novel interaction observed for NS1 and DIDO1 (Death Inducer-Obliterator 1), we further explored the role of DIDO1 in viral replication. Interactions between NS1 and DIDO1were corroborated in infected C6/36 and Aag2 cells, by colocalization and proximity ligation assays. Silencing DIDO1 expression in C6/36 and Aag2 cells results in a significant reduction in DENV and ZIKV replication and progeny production. Comparison of transcription analysis of mock or DENV infected C6/36 silenced for DIDO1, revealed variations in multiple gene expression pathways, including pathways associated with DENV infection such as RNA surveillance, IMD and Toll. These results suggest that DIDO1 is a host factor involved in the negative modulation of the antiviral response and necessary for flavivirus replication. Our findings uncover novel mechanisms of NS1 to promote DENV and ZIKV replication and add to the understanding of NS1 as a multifunctional protein.

**ABSTRACT IMPORTANCE:** Dengue is the most important mosquito-borne viral disease to humans. Dengue virus NS1 is a multifunctional protein essential for replication and modulation of innate immunity. To gain insights into NS1 functions, the protein interactome of dengue virus NS1 in Aedes albopictus cells was investigated using a proximity biotinylation system and mass spectrometry. Several protein pathways, not previously observed in vertebrate cells, such as epigenetic regulation and RNA silencing, were found as part of the NS1 interactome in mosquito cells. Among those, DIDO1 was found to be a necessary host factor for dengue and Zika virus replication in vertebrate and mosquito cells. Transcription analysis of infected mosquito cells silenced for DIDO1, revealed alterations of the IMD and Toll pathways, part of the antiviral response in mosquitoes. The results suggest that DIDO1 is a host factor involved in modulation of the antiviral response and necessary for flavivirus replication.

## INTRODUCTION

Mosquito borne flaviviruses, such as dengue virus, Zika virus, Japanese encephalitis virus, West Nile virus and Yellow fever virus represent a public health problem for the tropical and subtropical regions of the planet (1). Being human infectious viruses transmitted by vectors, these viruses need to adapt and take advantage of the cellular machinery of both the human host and their mosquito vector, to successfully complete their life cycle (2). The *Flavivirus* genome organization consists in one open reading frame encoding 3 structural (C (capsid), prM (precursor membrane), and E (envelope) and seven nonstructural (NS1, NS2a, NS2b, NS3, NS4a, NS4b, and NS5) proteins. The NS proteins are primarily responsible for viral replication and host immune evasion. The generated flavivirus polyprotein is processed co- and post translationally by the NS2b-NS3 protease complex; while the NS3-NS4a carry out helicase activities coupled with the NS5 which is responsible for capping, methylation, and replication of the viral genome (3–9). The small NS proteins, NS2A, NS2B, NS4A and NS4B are integral membrane proteins which do not have a known enzymatic activity (10). For DENV, the small NS proteins also have non-canonical roles which include acting as scaffolds and recruiters for replication complex formation, reorganization of cellular internal membranes, evasion from host immunity and metabolic changes during infection(10).

DENV NS1 is a glycoprotein of approximately 45-50 kD, that rapidly dimerizes after proteolytic maturation. Dimeric NS1 is located in the lumen of the ER and forms part of the replication complex as a scaffold protein (11, 12). NS1 also appears to be necessary for virion morphogenesis (13–15). In addition, NS1 is secreted from infected cells as a hexamer, and circulates in patients’ sera during the acute phase of viral disease (16, 17). Circulating NS1 have been associated with dengue pathogenesis by several mechanisms (12, 18). NS1 antigenemia in infected hosts promotes ZIKV infectivity and prevalence in mosquitoes(19). A recent protein interactome for DENV NS1 obtained in three different vertebrate cells lines showed that DENV NS1 interacts with at least 16 protein groups, as determined by gene ontology, including groups as diverse as mitochondrial proteins and proteins of the nuclear pore complex (20). Thus, flavivirus NS1 is a multifunctional protein whose functions are not yet fully understood (12, 21, 22).

Because NS1 is a ubiquitous protein among mosquito borne *Flaviviruses*, the functions and interactions described today for NS1 are the product of a collection of data from studies carried out using different members of the *Flavivirus* genus (23), as well as both vertebrate and mosquito cells. Although preserved functions for NS1 in both the human host and the vector mosquito are expected, recent evidence suggests that important differences in NS1 biology between vertebrate and mosquito cells also exist. For example, in mosquito cells NS1 has been reported to be translocated to the cell nuclei (24, 25), to lack a GPI tail, and to be absent from the plasma membrane; moreover, the traffic routes for secreted NS1 have been found to differ considerably between mosquito and vertebrate cells (6, 8, 9).

To gain information about the function of DENV NS1 in the mosquito, we implemented a proximity-dependent, biotin identification (BioID) assay to identify host proteins associated with NS1 in C6/36 cells (28, 29). This approach provides several advantages over other existing methods, including preservation of the cell architecture and allowed the derivation of a validated interactome of NS1 in live cells. Combining our interacting proteomic information with siRNAs and RNAseq assays, the putative epigenetic regulator DIDO1 was identified as an important host susceptibility factor for DENV and ZIKV infection in mosquito cells.

## RESULTS

### BioID in mosquito cells

We constructed four plasmids using conventional molecular cloning methods. As backbone, to obtain protein expression in mosquito cells, we choose the pAc5-STABLE1-Neo vector(30). We design 2 BirA-NS1 fusion proteins, with BirA located in the C- (Ac5-NS1-BirA-HA) or N-terminus (Ac5-Myc-BirA-NS1) of NS1 (Figure 1A). Each insert was cloned in frame upstream to GFP and the NeoR genes. The same plasmid containing only DEN2-NS1 sequence cloned in *Kpn*I-*Not*I sites and an empty plasmid Ac5-BirA-HA were used as controls. Western blot analysis of mosquito cells transfected with BirA-NS1 or NS1-BirA constructions showed the presence of NS1 with migration patterns compatible with NS1 fused with BirA (∼80KDa) or alone (∼48KDa). (Figure 1B). The reasons for the detection of non-fused NS1 is unknown since no additional proteolytic sites were found in these plasmids. Immunofluorescent assays showed expression of BirA as well as BirA-NS1 fused protein in transfected cells (Figure 1C). Finally, to demonstrate BirA activity in transfected C6/36 cells, cell lysates were analyzed by western blot and probed with streptavidin conjugated to horseradish peroxidase (HRP). Several different and discrete bands of varied intensities, and absent in non-transfected cells, indicated the presence of biotinylated proteins in the transfected cells (Figure 1D). These results indicated that mosquito cells were efficiently transfected and expressed NS1 fused to a functional BirA.

**Figure 1.**
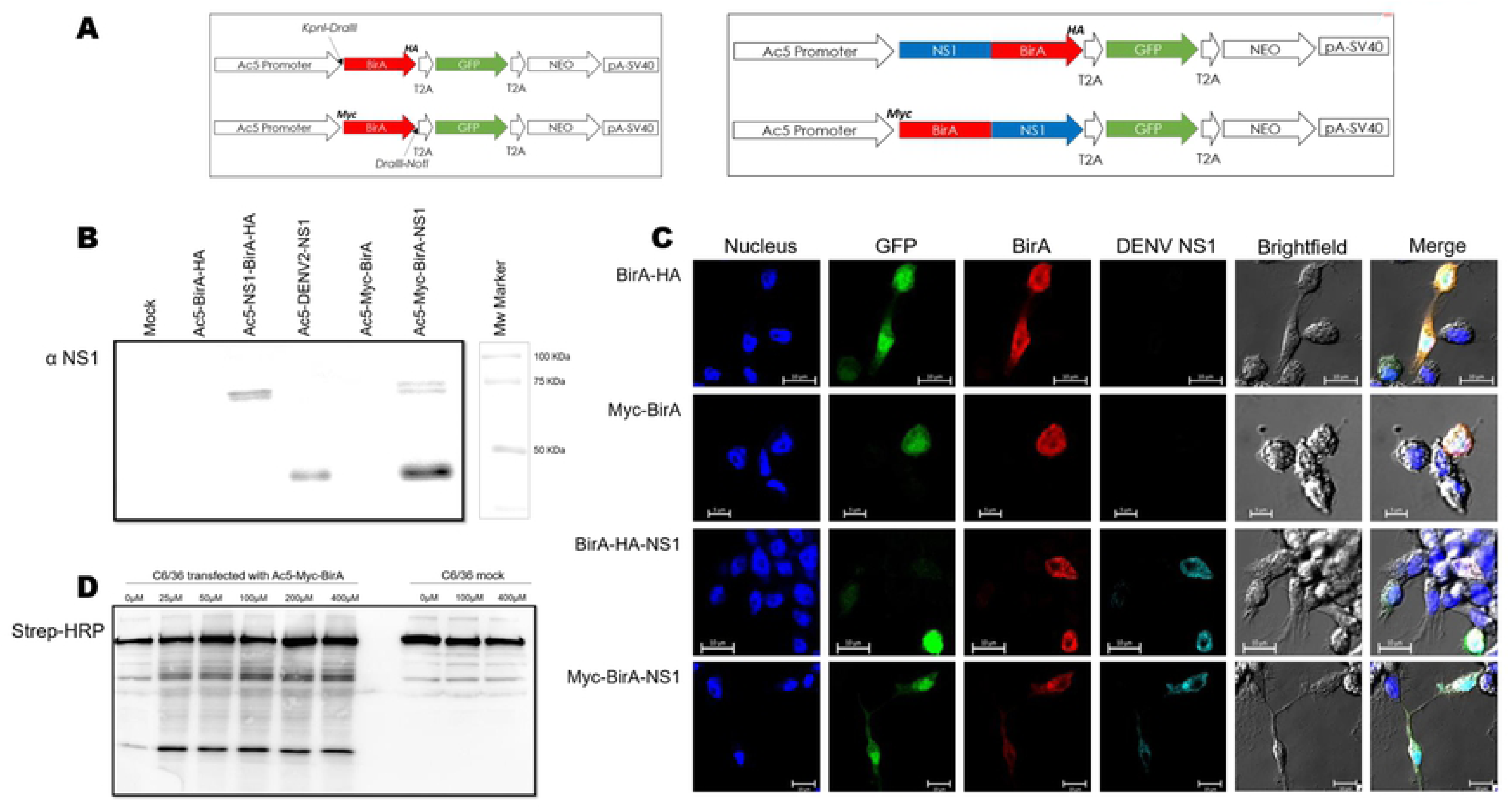
BioID expression system in mosquito cells. **(A)** Scheme of plasmids design used for NS1 protein interaction identification. Cloning sites for DENV2-NS1 are marked with arrows. For NS1-BirA fusion a HA flag and for BirA-NS1 fusion a Myc flag are detailed. T2A are self-cleaving peptides derived from thosea asigna virus. On left: empty vector and on right: fused genes **(B)** Western blot analysis of C6/36 cells transfected with plasmids carrying controls BirA-HA, Myc-BirA or DENV-NS1 alone or fused genes NS1-BirA-HA and Myc-BirA-NS1. Antibody anti NS1 were used for WB analysis. Mock: no transfected cells. **(C)** Confocal microscopy of C6/36 cells transfected with each plasmid. Transfected cells express GFP constitutively and nuclei were stained with DAPI, and cells immunostained with antibodies anti NS1 and HA and Myc flags. **(D)** Western blot analysis of C6/36 cells transfected with Ac5-BirA-HA. 24 hours after transfection cells were supplemented with biotin at concentrations indicated in the figure for 18 hours. Streptavidin-HRP were used to reveal biotinylated proteins.

### NS1 protein-protein interactions in mosquito cells

To identify mosquito cell proteins that interact with NS1, precipitated biotinylated proteins from mosquito C6/36 cells transfected with the experimental and control plasmids were identified by mass spectrometry. The overall average false discovery rate (FDR) was estimated (spectrum-level in raw data) to be 0.46 for true proteins. Only proteins with at least 2 unique peptides and more than 6 spectra were considered for further analysis. Proteins present in either NS1-BirA construction, but also present in BirA alone transfected cells, 197 in total, were excluded as non-informative, assuming that the tag was due to interactions with BirA itself (Figure 2). A total of 817 biotinylated proteins interacting with DENV-NS1 were found; 760 from cells transfected with the BirA-NS1 construction, 25 from cells with the NS1-BirA and 32 that were found in both conditions (**excel file1**). For literature validation, the protein interactions found here were compared with other NS1 protein interactions reported in the literature, regardless of the methodology or cell lines used (14–26) (**excel file2**). To compare the compilation of NS1 interactions reported in vertebrate cells with our results obtained in mosquito cells, a local BLAST to identify possible orthologues to each of the identified mosquito proteins was performed using NCBI human Refseq. A 724 of the proteins identified in mosquito cells had a match with the human database, while the rest (over 90) had no relevant equivalents, are hypothetical or uncharacterized proteins. To gain information about the functional enrichment and to construct functional ontology groups, an interactome analysis map was constructed using GeneMania (42) and String (43) plugins in Cytoscape (44) using the gene name annotations of these 724 proteins (Figure 2). Enrichment analysis showed multiple and diverse ontology groups, many were also found in vertebrate cells, including vesicle transport, the OST complex, CCT, translation, proteasome and metabolic processes, nuclear transport, ribosome, RNA modification and cytoskeleton (20, 32, 35, 37, 41). The high number of matches between the protein interactome obtained for DENV-NS1 in C3/36 cells with those reported in the mammalian cell literature and the coincidences among several functional ontology groups, indicates that the BioID assay carried out here is a reliable and effective method for the identification of protein-protein interactions.

**Figure 2.**
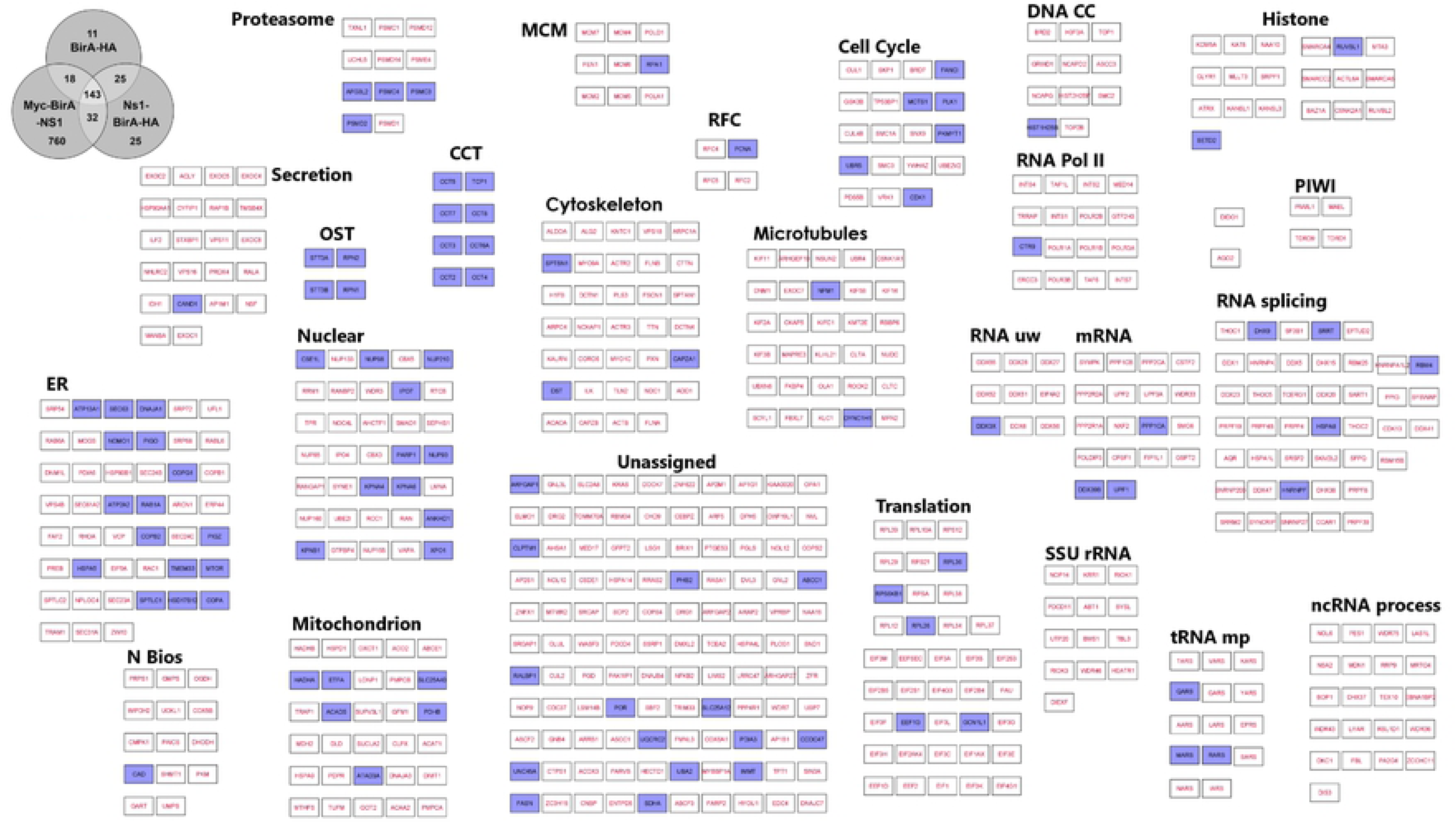
DENV2–NS1 protein interactions in mosquito cells. A total of 817 proteins were identified after comparative analysis with controls, discarding those also found in cells transfected with BirA alone. In the Venn diagram numbers represent the total of proteins identified for each condition. Because little information is available for mosquito proteins, an informative enrichment analysis was elaborated using human a protein database. Using local BLAST 780 human proteins functionally annotated were identified. These were used to construct an interactome analysis where proteins were clustered into functional groups using enriched GO terms, KEGG pathways and literature as a guideline. Highlighted in purple are proteins previously reported. Graphics analysis was performed with the program Cytoscape 3.8.2 using the String plugin. Abbreviations: **CCT:** chaperonin-containing T-complex GO:0005832; **Cell cycle:** KEGG pathway hsa04110; **Cytoskeleton:** GO:0005856; **Microtubules:** (cytoskeleton) GO:0015630; **DNA CC:** DNA conformation change GO:0071103; **ER:** endoplasmic reticulum GO:0005783; **Histone:** histone modification GO:0016570; **PIWI**: Piwi interacting RNA (piRNA) biogenesis Reactome Pathways HSA:5601884; **MCM:** minichromosome maintenance complex GO:0042555; **Mitochondrion:** GO:0005739; **mRNA:** mRNA surveillance KEGG pathway: hsa03015; **N Bios:** nucleoside monophosphate biosynthetic process GO:0009124; **ncRNA:** non-coding RNA processing GO:0034470; **Nuclear:** nuclear Envelope GO:0005635; **OST:** oligosaccharyltransferase complex GO:0008250; **RFC:** DNA replication factor C complex GO:0005663; **RNA Pol II:** GO:0016591; **RNA splicing:** GO:0008380; **RNA uw:** RNA secondary structure unwinding GO:0010501; **Proteasome:** GO:0000502; **Secretion:** GO:0046903; **SSU rRNA:** maturation of Small SubUnit ribosomal RNA GO:0030490; **Translation:** GO:0006412; **tRNA mp:** tRNA metabolic process GO:0006399.

Interestingly, other protein pathways related to epigenetic regulation (Histones, DNACC), RNA silencing (PIWI, ssuRNA, ncRNA processes), not previously reported in vertebrate cells, were also found as part of the NS1 protein interactome in mosquito cells (Figure 2). Among the novel proteins not previously reported as part of the interactome of NS1 in mammalian cells (36), and giving an strong signal in the NS1 interactome was the protein DIDO1 (Death-inducer obliterator 1 protein); a cytoplasmic protein that functions as a transcriptional regulator, and which has been found involved in apoptotic processes and embryonic stem cell development (45–47).

### Validation of NS1 interaction with DIDO1 in mosquito cells infected with DENV

The interaction between NS1 and DIDO1 was validated in DENV infected mosquito cells using confocal microscopy and proximity ligation assays (PLA, a technique capable of detecting direct protein-protein interactions in intact cells). Interactions between NS1 and the chaperon GRP78 (HSP5A) were also evaluated in parallel to serve as a positive control (38). No changes in DIDO1 distribution were observed after infection. Strong colocalizations (Pearson correlation coefficients above 0.6) were observed for NS1 and DIDO1 and GRP78 in DENV infected C6/36 cells fixed at 24 hpi, using confocal microscopy (Figure 3A). These results were confirmed using PLA in infected C6/36 cells, as well as in an additional mosquito cells line, Aag2, derived from *Aedes aegypti*. Clear PLA signals were observed for both proteins in C6/36 and Aag2 infected DENV2 cells fixed at 24hpi (Figure 3B). In addition, interactions between the NS1 from an additional DENV serotype, serotype 4 and another flavivirus, ZIKV, and DIDO1 were also tested by PLA and found to be strongly positive (Figure 3A and B). These results indicate that DIDO1 interact with NS1 in flavivirus infected mosquito cells.

**Figure 3.**
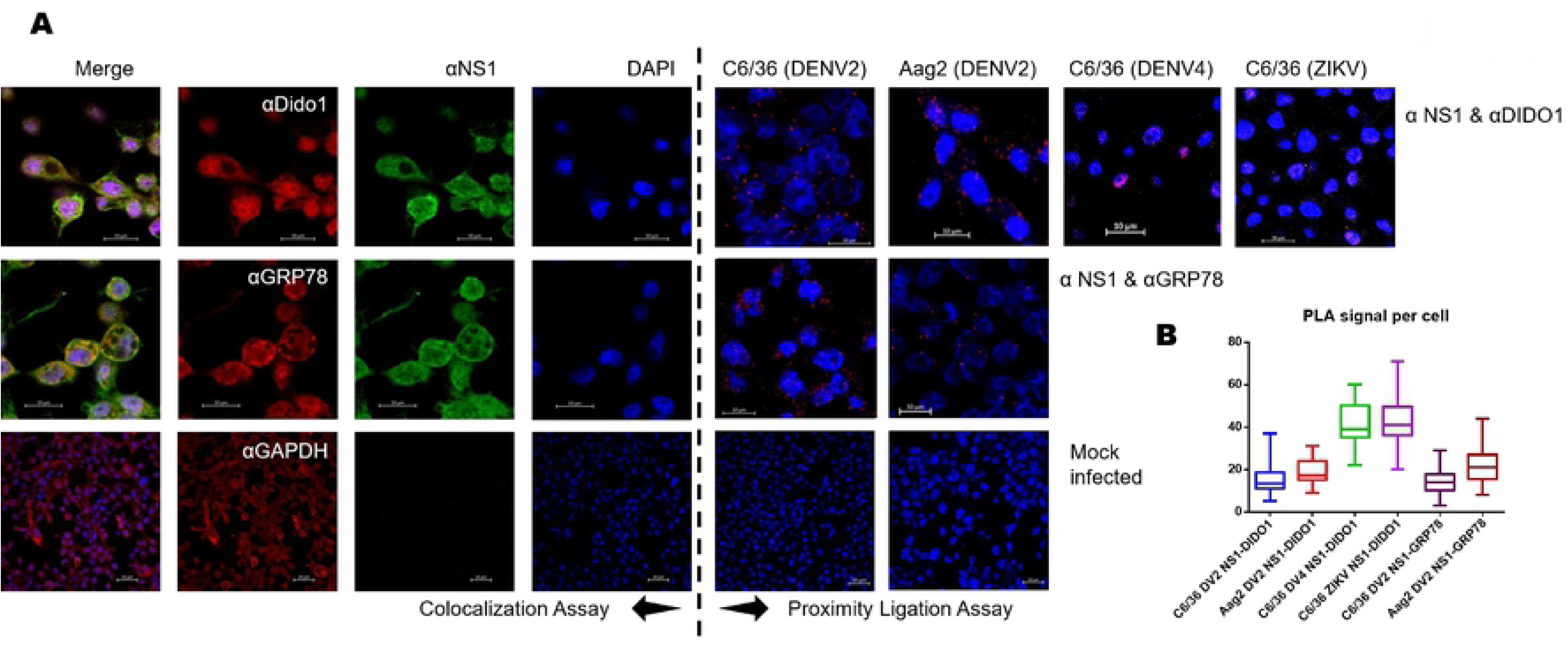
Validation of NS1 interactions with GRP78 and DIDO1. (A) Co-localization analysis of DENV-NS1 with target proteins. Left of dash line: Confocal microscopy was performed in C6/36 cells 48 hpi with DENV2. Immunostaining was performed using specific antibodies for each target. Anti-NS1 is shown in green, cellular proteins (GRP78 or DIDO1) are shown in red and DAPI is shown in blue. Right of dash line: Proximity ligation assay (PLA), of mosquito cells fixed at 48 hpi and performed following the manufacturer’s instructions. Red dots reveal positive proximity between NS1 and targeted mosquito protein. DENV2-NS1 PLA was performed for GRP78, and DIDO1 in C6/36 and Aag2 cells, and also for DENV4-NS1 and ZIKV-NS1 with DIDO1 in C6/36 cells. Experiments were carried out 3 times and typical results are shown. Non-infected cells used as negative controls are shown at the bottom (Mock). Scale bars represent 10 μm. (B) Colocalization analysis, Pearson correlation coefficients (PCC) was used to evaluate the degree of colocalization between NS1 and selected mosquito proteins. Bars represent means and ± standard errors. (C) PLA signal per cell quantifications. Signals of PLA were counted in maximum projection images for each assay, at least 30 cells were evaluated for each condition. Lines define minimum and maximum values and boxes represent standard errors divided by mean line.

### DIDO1 is a host susceptibility factor in mosquito cells infected with flavivirus

In silico analysis showed 38% homology between human and mosquito DIDO1 proteins (Figure 4A and B). Both proteins contain the 3 ordered domains (PHD, TFIIS and SPOC) compatible with the expected role of DIDO1 as a transcription factor. Although the mosquito protein does not show the splicing factor domain (SFD) at the C-terminus, it was possible to observe a similar sequence with many basic amino acids in the intermediate mosquito protein region (Figure 4B). A difference is the lack of exon 16 in mosquito DIDO1 which has been implicated in protein function regulation (48); in contrast, the annotated mosquito protein includes a large amino terminal sequence. Another difference is the presence of two exclusive domains, SWIRM in human DIDO1 and BRK in mosquito DIDO1, which are both also associated with transcription. Thus, despite some differences in structure, multiple similar functions can be expected for DIDO1 in human and mosquito cells.

**Figure 4.**
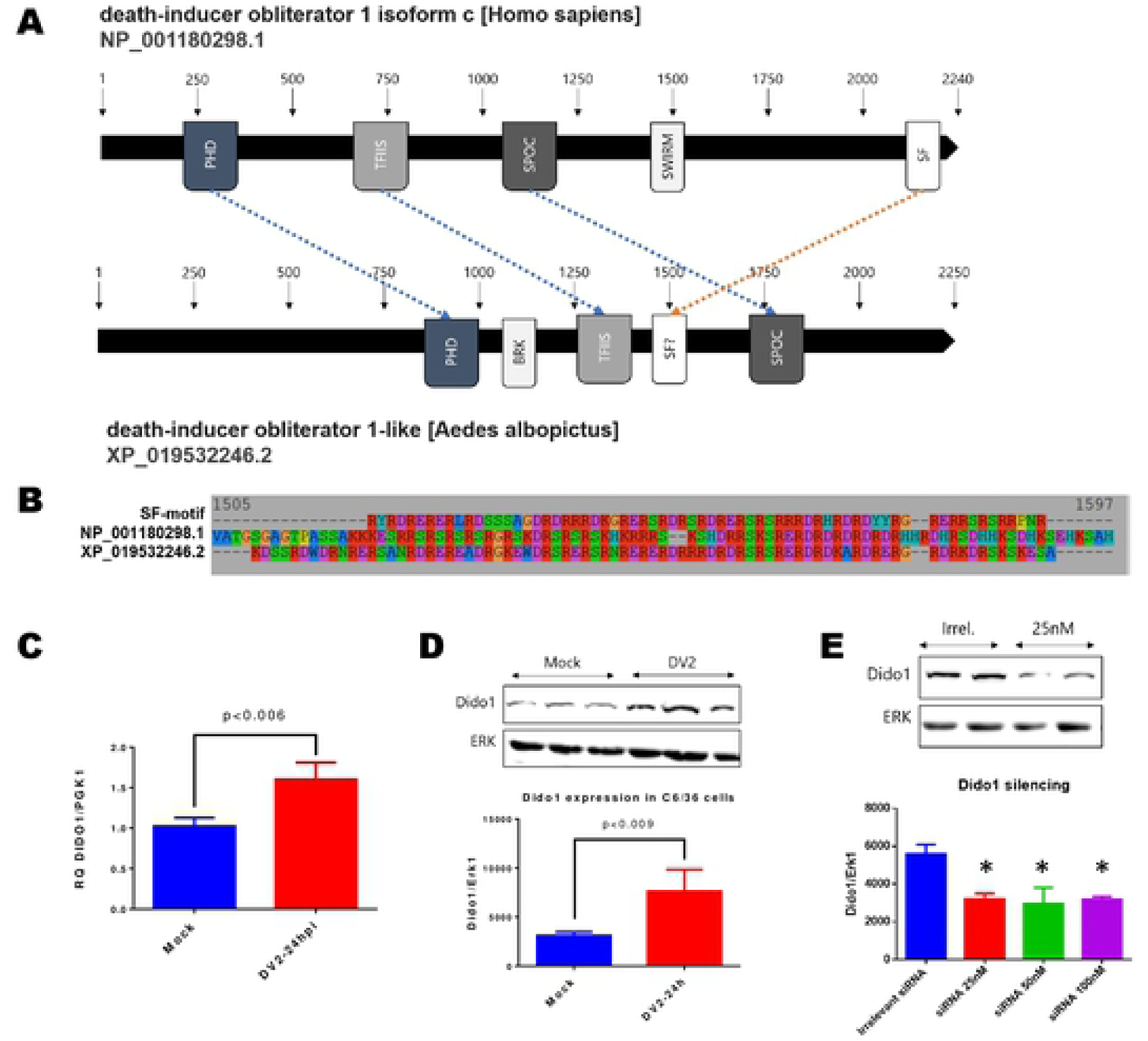
Death-inducer obliterator 1 in mosquito. (A) Schematic representation of domain organization of *Aedes* Dido1 protein compared to human orthologue, showing 38% homology between amino acid sequences. *Aedes* DIDO1 has at least 3 domains in the same order (PHD, TFIIS and SPOC domains). Domains SWIRM and SF are not found in mosquitos, which also have a differential BRK domain. A similar sequence of SF domain in *Aedes* Dido1 sequence, characterized by having atypical number of basic amino acids. (B) protein sequence alignment of putative DIDO1 SF domain. (C) mRNA levels of DIDO1 increased relative to PGK1 in C6/36 cells infected with DENV2 at m.o.i. of 3, 24 hours post infection. Difference was statistically significant. (D) DIDO1 protein expression increased in mosquito cells 24 after infection with DENV2. Western blot analysis shows more expression of Dido1 relative to ERK expression (upper panel), the difference between Mock and infected cells is statistically significant (bottom panel). (E) Dido1 knockdown using commercial siRNAs. Western blot analysis shows nearly 50% diminished expression of DIDO1 relative to ERK expression (upper panel) using siRNA at 25nM or higher concentration. Assays were performed by triplicate and bars represent means and ± standard errors. *=*p≤0.005*.

Messenger RNA levels as well as levels protein expression for DIDO1 were found significantly increased at 24 hpi in DENV infected cells, indicating a possible demand during viral replication (Figure 4C and D). To probe this hypothesis, DIDO1 silencing was standardized to evaluate the effect of lack of this protein in flavivirus replication. As shown in Figure 4D, up to 50% reduction in DIDO1 expression was achieved when 25 nM siRNA final concentration was used, and not further reduction was observed at higher concentrations. Nonetheless, for consistency 50nM final concentration of DIDO1 siRNA was chosen to perform the infection assays. This concentration was not found to cause cytotoxicity in either C6/36 or Aag2 cells after 48 h of transfections, as indicated by MTT assays (Figure 5). DIDO1 knockdown resulted in a significant decrease of viral progeny (up to 1,5 logs) in C6/36 and Aag2 cells infected with DENV2 (Figure 5A). The negative effect of DIDO1 silencing in DENV replication was also observed in DENV serotype 4 infected cells. In addition, ZIKV yield was affected by nearly 1 log in mosquito cells silenced for DIDO1 (Figure 5A). Genome quantification by qRT-PCR indicated that the number of viral genome copies of DENV2 and ZIKV was also decreased in DIDO1 silenced cells (Figure 5B); as well as the expression of viral proteins NS1 and prM in cells infected with DENV2 (Figure 5C). These results taken together indicate that DIDO1 is an important host susceptibility factor for DENV and ZIKV replication in mosquito cells.

**Figure 5.**
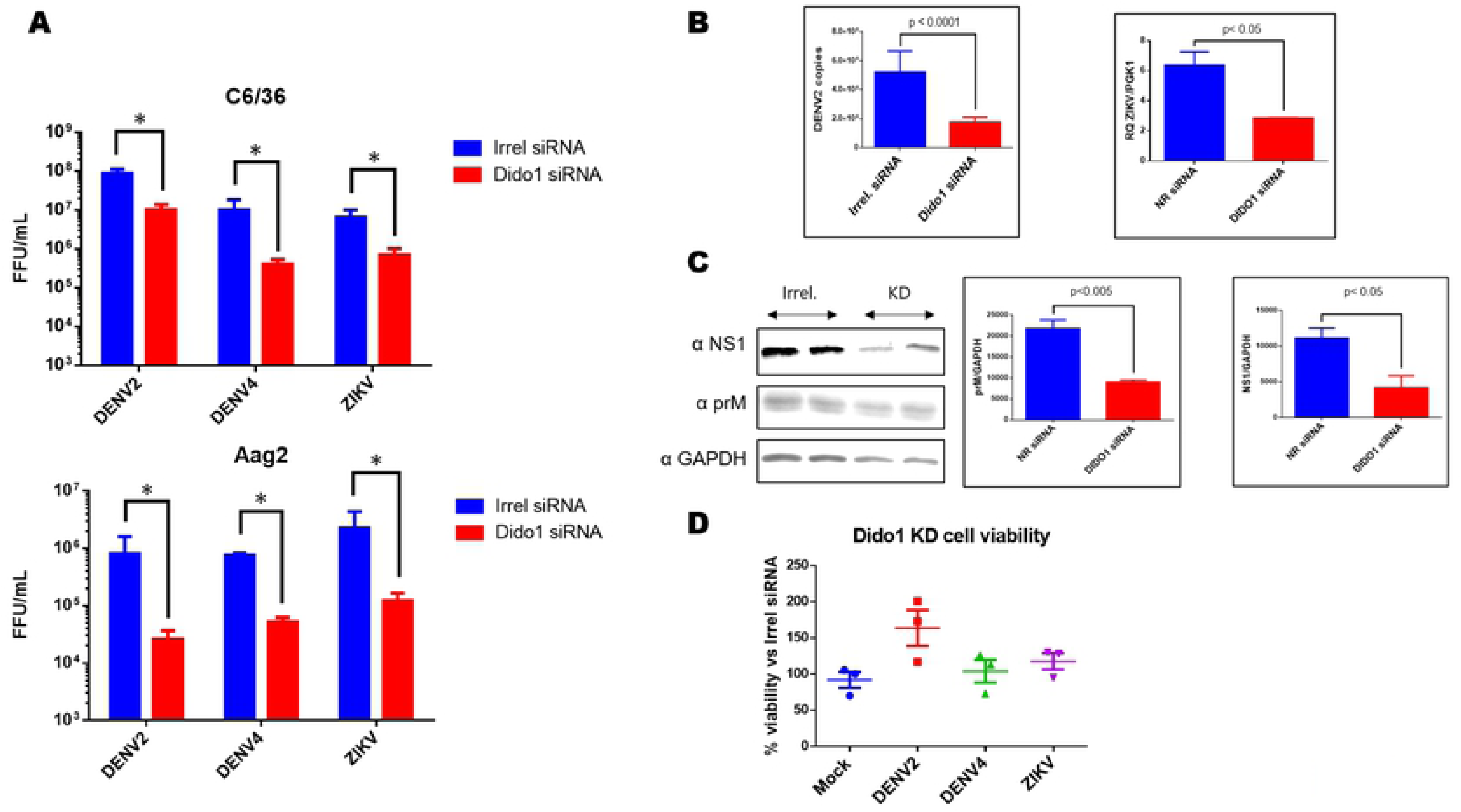
DIDO1 is a flavivirus host susceptibility factor in mosquito cells. (A) Viral progeny of DENV2, DENV4 and ZIKV in C6/36 and Aag2 cells silenced for DIDO1 expression, as assayed by focus forming units in cell supernatants collected 24 hpi. (B) Viral RNA measured by qPCR in C6/36 cells infected with DENV2 or ZIKV and silenced for DIDO1, harvested at 24 hpi (C) Western Blot of DENV2 viral proteins NS1 and prM assayed in lysates silenced for DIDO1 and harvested 24 hpi (D) Cell viability in mock or infected cells silenced for DIDO1 and assayed by MTT assays 24 hpi. All experiments were carried out by triplicate, bars represent means and ± standard errors. *=*p≤0.005*.

Finally, to test if DIDO1 is also a necessary protein for DENV replication in vertebrate cells, BHK21 cells were silenced and infected with DENV. As shown in Figure 6, DIDO1 siRNA treatment of BHK-21 cells reduced DENV yield in nearly 2 logs (Figure 6C), as well as the expression of viral proteins NS1 cells infected with DENV2 (Figure 6B). These results suggest that DIDO1 is also a host susceptibility factor for DENV in vertebrate cells.

**Figure 6.**
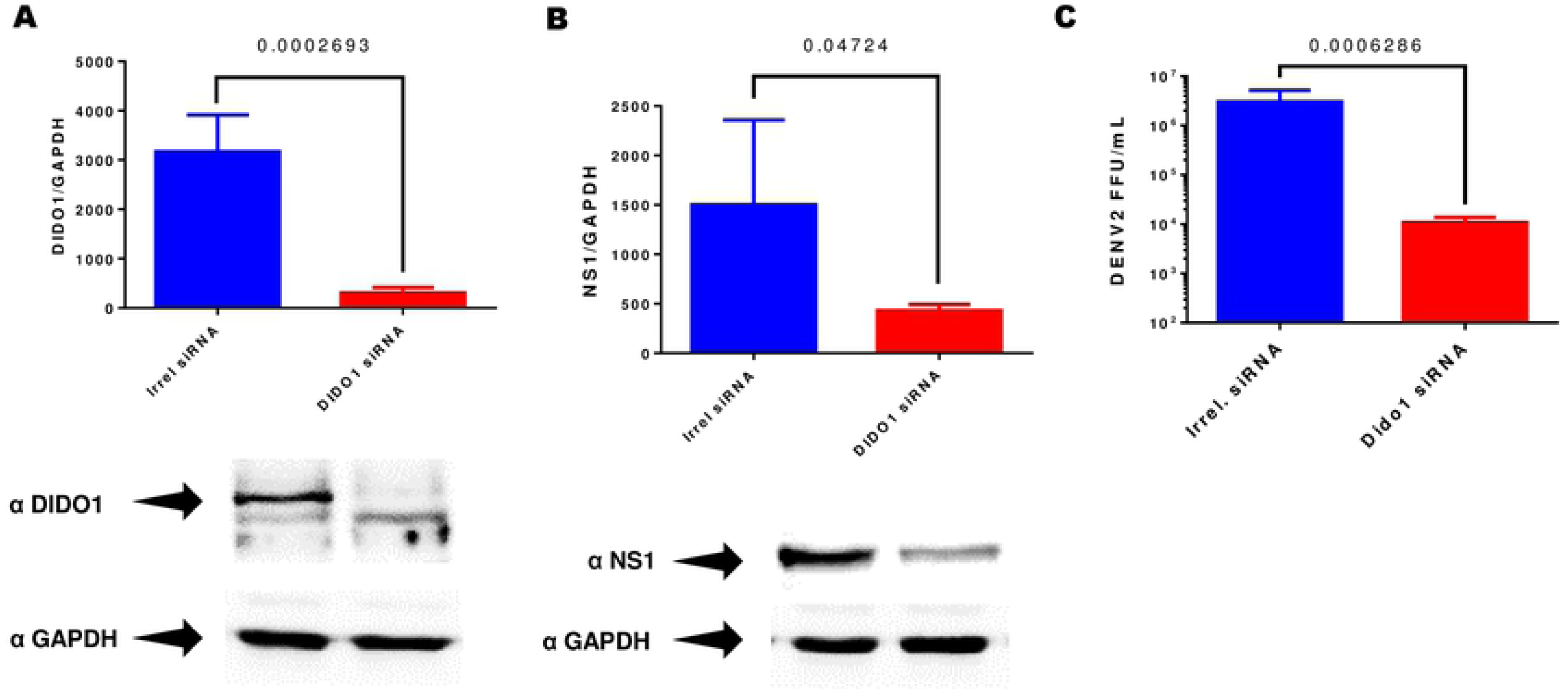
Silencing of DIDO1 in BHK21. (A) Western blot analysis shows nearly 90% diminished expression of DIDO1 relative to GAPDH expression (lower panel) using DIDO siRNA at 50nM. (B) Western Blot of DENV2 viral proteins NS1 assayed in lysates silenced for DIDO1 and harvested 24 hpi. (C) Viral progeny of DENV2 in C6/36 cells silenced for DIDO1 expression assayed by focus forming units in cell supernatants collected 24 hpi. Assays were performed by triplicate and bars represent means and ± standard errors. P value is indicated.

### Transcriptomic analysis in infected C6/36 cells silenced for DIDO1

To gain information about the functional relationship between DIDO1 and DENV NS1 during viral infection in mosquito cells, RNA sequencing was carried out in DENV infected C6/36 cells transfected with siRNA specific for DIDO1 or an irrelevant siRNA, as a control. In addition, RNA sequencing analysis were also carried out in mock infected cells also silenced or not for DIDO1. For each of the 4 conditions, more than 70M reads were obtained and mapped to the annotated genome of *Aedes albopictus* in the NCBI database. All samples had similar overall results with more than 50% of reads that matched to 18,699 genes and 29,746 different transcripts (excel file3). To better understand the differential expression gene (DEG) analysis, 3 combinations were compared: mock infected C6/36 cells silenced or not for DIDO1; C6/36 cells infected or not with DENV2 and finally DENV infected C6/36 cells silenced or not for DIDO1 (Figure 7). Using a cutoff p-value < 0.05, a log2 fold-change (log2FC) expression ±2 was observed for an important number of genes in all 3 combinations compared (Supplemental Figure 1). To obtain functional information of DEGs in each comparison, gene ontology (GO) analysis was carried out using the ClueGO (49) plugin in Cytoscape (44). Due to the lack of annotations in the genome of *Aedes albopictus*, *Aedes aegypti* homologues were used for the GO analysis. Bar plots in Figure 7 detail the main findings derived from GO of DEG. In agreement with previous data obtained in C6/36 cells infected with ZIKV (50) or Aag2 cells infected with DENV2 (51), DENV infected cells showed alterations in multiple pathways related to biosynthesis and energy production, in comparison with non-infected cells (Figure 7), and thus validating our preliminary analysis of DEG. In the case of C6/36 mock infected cells treated with DIDO1 siRNA versus the irrelevant siRNA used as a control, GO of DEG shows multiple pathways associated to cell cycle regulation and differentiation such as Notch, MAPK, FoxO, Hedgehog, mTOR, TGF-beta, Wnt and Hippo pathways; in addition, changes in gene expression of proteins related to autophagy and apoptosis, not found in the comparison between infected and not infected cells, were also found (Figure 7). It has been reported that DIDO1 is a switchboard that regulates self-renewal and differentiation invertebrate embryonic stem cells (46) and probably participates in apoptosis (52) which is consistent with the changes in gene expression observed in mock infected C6/36 silenced for DIDO1, and suggests conservation of functions of DIDO1 between vertebrate and mosquito cells. Notably, in DENV infected cells silenced or not for DIDO1, modifications in the expression of genes associated with autophagy, apoptosis, and the Toll and IMD pathways, which in invertebrates activate the *Relish* (NF-κB) transcription factor, were observed. The RNA seq results obtained in silenced, infected cells suggest that DIDO1 may participate in the modulation of the innate immunity and antiviral response in the mosquito cell.

**Figure 7.**
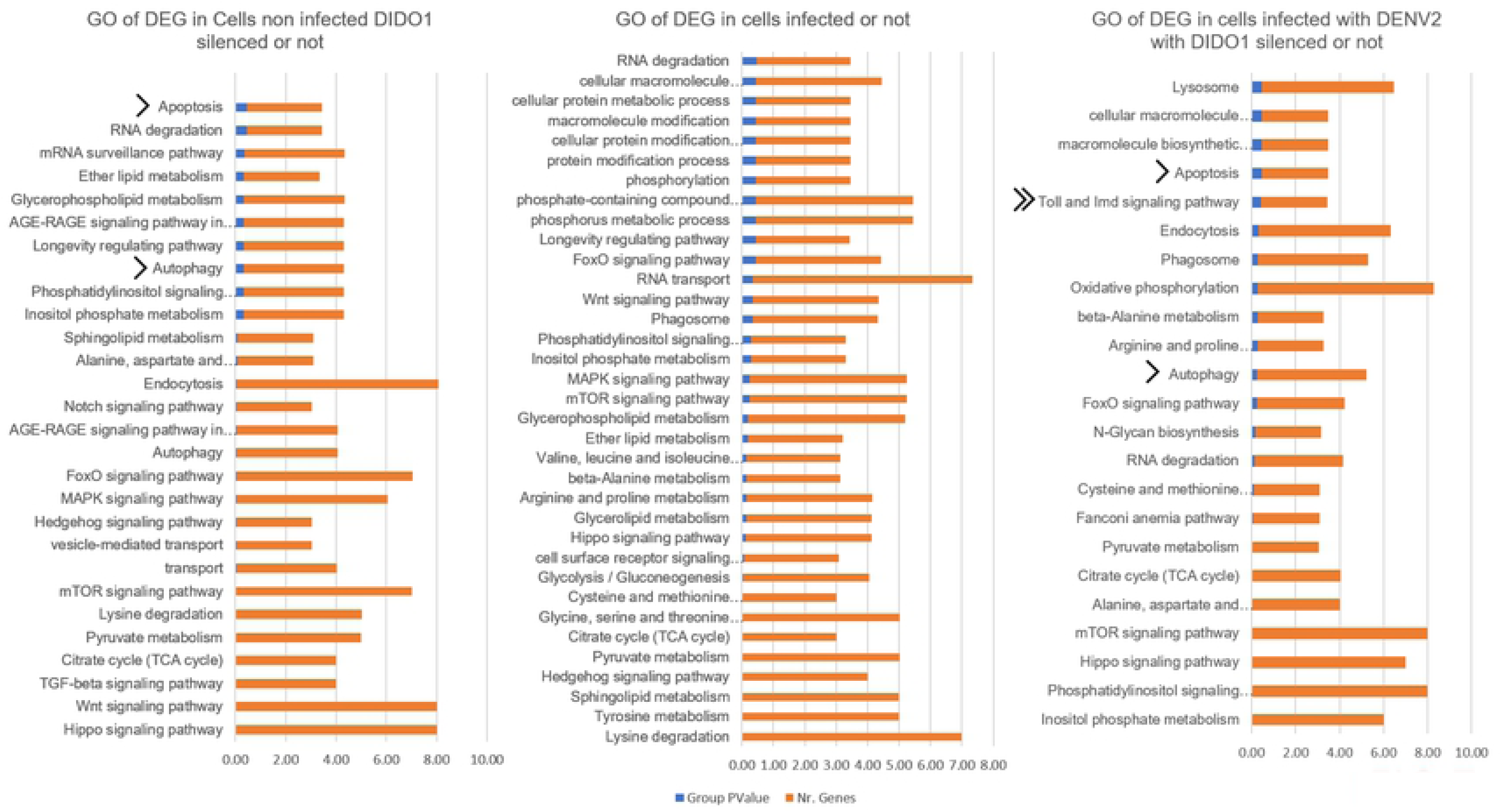
C6/36 cells transcriptomic analysis under DIDO1 knockdown and/or infection with DENV2. Gene ontology analysis of DEG. It was considered cell component, biological process, and KEGG pathways proteins annotations with p-value < 0.5 for groups. Due to the lack of functional information for *Aedes albopictus*, analysis was derived from *Aedes aegypti* protein homologues.

Finally, DEG analysis with DIDO1 silenced samples, independently of the infection status, resulted in 9 top genes affected (Supplemental Figure 2), besides 3 uncharacterized genes, the rest of them are all functionally related to RNA modification, in agreement with the recent findings pointing to DIDO1 as a participant in spliceosome and transcriptional regulation (48).

## DISCUSSION

The DENV NS1 protein is a multifunctional protein that is located in the replication complexes and is essential for viral replication. Yet, in addition to the scaffold function as part of the replication complexes, little is known about the mechanisms that make DENV NS1 as essential viral protein for replication. NS1 is present and highly conserved in all mosquito borne flaviviruses and defining a global interactome for NS1 protein has been difficult due to the widely different experimental approaches found in literature. A recent global proteomic analysis of DENV NS1, in combination with a functional RNAi screen, allowed the identification of 270 host proteins that interact with NS1 (20). The results showed that NS1 is involved with multiple cell organelles, and signaling pathways, beyond the expected ER and Golgi complex, as for example components of the nuclear pore (XPO5, TNPO1) and the machinery of DNA repair and replication (FANCI, PCNA, MSH2, and MCM3), supporting the predicted multifunctional nature of NS1. Thus, to gain knowledge about the functions of NS1 in the infected cell, particularly the mosquito cell, where such studies are scarce, we set out to identify the protein interactions of DENV NS1 in C6/36 cells. To this aim, we first adapted the BioID system to function in mosquito cells. The system allowed us to identify a total of 817 proteins interacting with DENV NS1 in C6/38 cells. The number of protein and systems coincidences obtained with previous studies (bibliography validation) using very different experimental approaches, including yeast two-hybrids (31, 34, 40, 53, 54), immunoprecipitations (20, 32, 34, 35, 37, 38, 41), computational predictions (55) and structural inferred interactions (56). Moreover, experimental validation for 2 of proteins identified was obtained in DENV infected cells, adding confidence to our global NS1 interactome obtained in mosquito cells.

Using a basic comparative analysis by gene names, nearly 14% (100 proteins) of the interacting proteins matched previously reported for NS1 PPIs in vertebrate cells. Functional coincidences are notably higher including proteins from the oligosaccharide transferase complex (OST), the chaperonin containing TCP-1 (CCT), nuclear import and export, vesicle localization and ribosomal proteins. Interestingly, other proteins related with epigenetic regulation pathways such as histone reader and modifiers (KAT8, MLLT3, BRPF1, KDM5A, KANSL3, KANSL1, GLYR1 and NAA10) or part of SWI/SNF transcriptional complex (SMARCA4, SMARCC2, SMARCA5 and MTA3) and RNA silencing (AGO2, PIWIl1, MAEL, TDRD9 and TDRD1), not previously reported in vertebrate cells, were also found as part of the interactome of DENV NS1 in mosquito cells. The abundant presence of NS1 reported in the cell nuclei of DENV infected C6/36 (24, 25) and which is not so evident in infected vertebrate cells, may explain in part the novel associations uncovered for NS1 with epigenetic regulatory systems, nuclear or RNA splicing components. The role played by NS1 in cell nuclei is currently unknown, but functions in gene regulation of spliceosome modulation have been reported for other DENV proteins, such as NS5, which enter the cell nuclei during infection (57). Unfortunately, comparative analysis of human and mosquito NS1 interactomes are challenging, due to the lack of reliable annotations and validation of the mosquito databases and the intrinsic complexity and plasticity of the genome of this species, and further differences may be evident in the degree that mosquito gene databases are better cureated.

Changes in the expression mRNA and protein levels of DIDO1 in DENV infected cells suggested a demand for and role of DIDO1 during infection. DIDO1 is a switchboard, transcription factor that has been implicated in the regulation of apoptosis, cancer cell proliferation and maintenance of embryonic stem cells by complex regulatory loops (46–48, 58). In mosquito cells, the existence of splicing variants for DIDO1 is unknown but the *DIDO1 gene* and a protein orthologue for DIDO1 with conserved domains were clearly identified. DIDO1 knockdown resulted in a significant reduction in DENV and ZIKV viral replication, without any significant effect on cell survival. Reductions in viral progeny were accompanied by reductions in viral protein synthesis and viral genomic RNA copy number. These results suggest that DIDO1 is a host factor necessary for DENV and ZIKV replication in the mosquito vector. Interestingly, DIDO1 also seem to be a necessary host factor for DENV in vertebrate cells as suggested by the results obtained in BHK21 cells. The reasons why DIDO1 was not detected in previous studies as part of the NS1 interactome are unknown but may be related to the low levels of NS1 in the nuclei of infected vertebrate cells.

No previous participation of DIDO1 in any viral cycle has been reported. Thus, a non-biased transcriptomic analysis using C6/36 cells silenced for DIDO1, infected or not with DENV2 was carried out, in an attempt to understand the mechanisms by which the interaction between DIDO1 and NS1 might favor DENV replication. DEG analysis revealed variations in multiple targets consistent with the molecular role of DIDO1, which could be related to transcription and/or epigenetic regulation. In infected cells silenced for the expression of DIDO1, DEG analysis indicated the alteration of the Toll and the IMD pathways, none of which were observed in infected cells transfected with the irrelRNA. Moreover, alterations of apoptosis and autophagy pathways were observed in either mock or infected cells silenced for DIDO1, which are both cell responses aimed to control viral infections, and in agreement with previous data pointing to DIDO1 as a regulatory factor for apoptosis. Thus, the DEG data suggest that the presence of DIDO1 is necessary to negatively modulate the activation of several antiviral response pathways during DENV infection. Evidence showing the capacity of DIDO1 to modulate RNA splicing in insects has been found (59). The viral targeting of key regulatory proteins implies that several major processes, relevant for the virus life cycle, may be modulated simultaneously.

In summary, the data taken together is consistent with the notion that DIDO1 interaction with NS1, results in diminished innate immune responses to promote viral replication in the mosquito cell. A gap exists in the understanding of the susceptibility host factors necessary for DENV and other flavivirus replication in the mosquito vector versus the vertebrate host. This gap needs to be closed if effective measurements to reduce the vector capacity of the mosquitoes are to be developed.

## MATERIALS AND METHODS

### Plasmid constructions

Plasmid Ac5-Stable2-neo was a kind gift from Rosa Barrio and James Sutherland (Addgene plasmid # 32426; http://n2t.net/addgene:32426; RRID:Addgene_32426) (30). Donor plasmids for BirA pcDNA3.1-MCS-BirA-HA and pcDNA3.1-Myc-BirA were a kind gift from Carlos Sandoval and Susana López (Instituto de Biotecnología-UNAM, Mexico). The NS1 protein of DENV serotype 2 was subcloned from a plasmid provided by Sebastián Aguirre y Ana Fernández Sesma (Icahn School of Medicine, Mount Sinai, New York). Cloning was carried out by gene amplification using primers with adapters for site restriction. For NS1 cloning in N terminal, the following primers were used: KpnI-BirA: ATAGCTAGGTACCGCCACCATGGAACAAAAACTCATCTC; BirA-DraIII-NotI: TCCTTCGCGGCCGCCATAGCTCACTACGTGGAGCTTCTCTGCGCTTCTC; DraIII-NS1: ATAGCTACACGTAGTGGCCACCATGGGATCACGCAGCACCTCACTGTCTGTG and NS1-NotI: ATCTTCGCGCGGCCGCCAGCTGTGACCAAGGAGTTGAC. And for NS1 cloning in C terminal, the following primers were used: KpnI-DraIII-BirA: ATCTTCGGTACCATAGCTCACGTAGTGGCCACCATGGGACGCTTAAGGCCTGTTAACCGGTCG TAC; BirA-NotI: TCCTTCGCGGCCGCCTGCGTAATCCGGTACATCGTAAG; KpnI-NS1: ATAGCTAGGTACCGCCACCATGGGATCACGCAGCACCTCACTGTCTGTG and NS1-DraIII: ATAGCTCACTACGTGAGCTGTGACCAAGGAGTTGAC.

First, Myc-BirA was amplified using primers KpnI-BirA / BirA-DraIII-NotI and cloned in Ac5-Stable2 eliminating the m-Cherry coding sequence by restriction with KpnI/NotI enzymes. Once the gene substitution was corroborated, the DENV2-NS1 gene was inserted. For this construction, the NS1 sequence was amplified with DraIII-NS1/NS1-NotI primers. Cloning was conducted by restriction with DraIII-NotI enzymes and ligation. In addition, BirA-HA was amplified using primers KpnI-DraIII-BirA / BirA-NotI and cloned in Ac5-Stable2 also eliminating m-Cherry coding sequence by restriction with KpnI/NotI enzymes. Then, the DENV2-NS1 sequence was amplified with KpnI-NS1 /NS1-DraIII primers and cloned by restriction with KpnI-DraIII enzymes and ligation. All 4 constructions were corroborated by restriction analysis and DNA sequencing; protein expression was corroborated by western blot using anti-NS1, anti-Myc or anti-HA antibodies and fluorescent microscopy.

### Cell lines

Mosquito cells C6/36 from *Aedes albopictus* (ATCC CRL-1660) and Aag-2 from *Aedes aegypti* (a kind gift by Fidel de la Cruz, CINVESTAV) were grown at 28°C with 5% CO_2_ in Eagle’s minimum essential medium (EMEM) supplemented with 5% fetal bovine serum (FBS) and 100 U/ml penicillin-streptomycin. Vertebrate cells BHK-21 (ATCC CCL-10) and Vero-E6 (ATCC CRL-1586) were grown at 37°C with 5% CO_2_ in EMEM supplemented with 5% or 10% FBS and 100 U/ml penicillin-streptomycin.

### Virus strains

Dengue virus serotype 2 (DENV2) strain New Guinea, dengue virus serotype 4 (DENV4) strain Philippines H241, were generously provided by Mauricio Vázquez (Laboratory of Arboviruses and Hemorrhagic Viruses, Institute of Epidemiological Diagnosis and Reference [InDRE], Mexico). Zika virus (ZIKV) strain MR77-Uganda was a kind gift by Susana López (IBT-UNAM). DENV strains were propagated in suckling mouse brains (ICR; CD-1); animals were provided by the Unit of Production and Experimentation of Laboratory Animals (UPEAL-CINVESTAV) and handled compliying with the ethical procedures of our Institution. The ZIKV strain was propagated in C6/36 cells. DENV and ZIKV were titrated by focus-forming assay in BHK-21 cells or Vero E6 cells, respectively (45). Briefly, serial dilutions of viral stocks or supernatants of experiments were diluted in serum-free medium and added to cell monolayers grown in 96-well plates. Virus absorption was allowed for 2 h at 37°C; then EMEM–10% FBS was added and incubated for an additional 48 h before fixation and quantitation. Infected cells were visualized by labeling with anti-flavivirus E-glycoprotein antibody using a mouse VECTASTAIN ABC-horseradish peroxidase (HRP) kit (PK-4002; Vector Laboratories) and a DAB peroxidase substrate kit (SK-4100; Vector Laboratories).

Virus infections of either C6/36 or Aag2 mosquito cells were carried out at a multiplicity of infection of 3 (MOI=3) and depending on the experimental needs, fixed at 24 or 48 hpi.

### Cell transfections

Plasmids were transfected into 85% confluent monolayers of C6/36 cells seeded in 24-well plates. Cells were transfected with 1 µg of plasmid DNA and 2 µL of Lipofectamine 2000 reagent (Invitrogen). Six hours post transfection, cells were supplemented with EMEM at 10% FBS. Antibiotic selection with G418 was not possible for transfected cells with NS1 containing plasmids due to apparent toxicity after a week of selection. Transfection efficiency was estimated at 50-60 % by fluorescence microscope and experiments were carried out at 24-48 hours post-transfection.

### Knockdown of DIDO1 gene expression

Cells were transfected with a mix of DIDO1 siRNAs (FlexiTube™ GeneSolution GS11083 for DIDO1, 1027416, Qiagen) using HiPerFect transfection reagent (Qiagen). AllStars™ negative-control siRNA (Qiagen) was used as an irrelevant siRNA control at the same concentrations as the gene specific siRNAs. Gene silencing experiments were carried out in cells grown in 24-well plates and silencing efficiency corroborated by western-blot using a commercial anti-DIDO1 antibody (cat. GTX 59722; GeneTex). Protein expression was normalized to the expression of glyceraldehyde-3-phosphate dehydrogenase (GAPDH) (GTX100118; GeneTex), or ERK-1 (sc-281291; Santa Cruz Biotechnology, Inc.). Expression levels relative to negative-control cells, transfected with the irrelevant siRNA, were estimated by densitometric analysis with ImageJ software (60).

### Quantitative real-time PCR for viral RNA

Levels of viral genomic RNA copies were quantified by real-time RT-PCR. After infection times, cell monolayers were washed, and the total RNA isolated with TRIzol reagent (Invitrogen) according to the manufacturer’s procedures. A total of 0.1µg of total RNA was used to determine the copy number of genomic DENV2 RNA using a standard curve with serial dilutions of viral stocks. For ZIKV RNA, copy number estimation was performed by relative quantification using the housekeeping PGK1 gene expression as reference (61). One step RT-PCR was carried out using the QuantiTect probe RT-PCR kit (Qiagen, Valencia, CA) with 500 nM forward and reverse primers and 50 nM labeled probes (DENV2-FAM and ZIKV-FAM and PGK1-VIC TaqMan). Detection primers and probes for DENV and ZIKV were as described by Chien et al. (62) and Lanciotti et al. la(63), respectively. Cycling conditions for ZIKV and PGK1 were 50°C for 30 min, 95°C for 15 min, 45 cycles of 95°C for 15 s, and 60°C for 1 min. Cycling conditions for DENV2/4 were 50°C for 30 min, 95°C for 15 min, 50°C for 30 s, 72°C for 1 min, 45 cycles of 95°C for 15 s, and 48°C for 3 min. Amplification was done in a StepOne real-time PCR device from Applied Biosystems (Applied Biosystems, Foster City, CA), and results were analyzed using StepOne software v2.3. Each assay was performed in three biological replicates with two technical replicates, and each assay included no-template negative controls and DENV2, DENV4, and ZIKV positive controls. Results were expressed as the total number of RNA copies or RQ (2^-(delta-delta Ct)^) using relative expression of viral genome versus PGK1 expression.

### Cell viability assays

Cell viability was determined with the Cell Titer 96 AQueous non-radioactive cell proliferation assay, used according to the manufacturer’s recommendations (MTS assay, G3580; Promega). Cells seeded in 96-well plates were treated with DIDO1 siRNA or irrelevant siRNA, both at 100nM final concentration, and kept for 48 hours after transfection. Viability was expressed as a percentage of the control condition, taken as 100% viability. Three biological replicates were used for each condition.

### Confocal microscopy assays

Confluent cell monolayers grown in 24-well plates containing glass coverslips, were infected with DENV or ZIKV with MOI of 3. Twenty four or 48 hours after infection, cells were fixed in 4% paraformaldehyde and permeabilized with 0.2% Triton X-100, incubating for 10 min at room temperature in each step. For immunostaining, Mab2B7 anti-NS1, a kind gift by Eva Harrys (University of California, Berkeley, USA) (64) and the commercials anti-DIDO1 (GTX59722), anti-GRP78 (GTX22902) anti-HA-tag (GTX30545), anti-myc-tag (GTX30518) antibodies (Genetex) were used at 1/300. Conjugated anti-mouse Alexa-488 or Alexa-598, anti-goat Alexa-568, and anti-rabbit Alexa-648 or Alexa-488 (donkey pre adsorbed secondary antibodies; Abcam) were used at 1/500. Nuclei were stained with DAPI. Coverslips were mounted in Fluoroshield™ with DAPI (Sigma-Aldrich). The images were analyzed using a Carl Zeiss LSM 700 confocal microscope. Pearson correlation coefficients (PCC) were obtained from at least 30 independent cells from confocal images to evaluate the colocalization between proteins of interest and NS1. Icy Image software was used for image analysis and PCC values calculations with colocalization studio plugin (65).

### Proximity ligation assays

To confirm the interaction between NS1 and GRP78 and DIDO1 proteins, we used the proximity ligation assay (PLA) Duolink PLA kit (Sigma-Aldrich). Mosquito cells C6/36 or Aag2 seeded over slides in 24 well format plates, were infected at MOI=3. Twenty four hours after infection, cells were processed as recommended by the manufacturer. In brief, cells were fixed in cold methanol, permeabilized and incubated with a blocking agent for 1 h. Samples were incubated overnight with commercial primary antibodies generated in rabbit for the cellular proteins or in mouse for the viral NS1 protein, used 1/100. Duolink PLA probes detecting rabbit or mouse antibodies were applied to the slides, and incubated for 1h in a preheated humidity chamber at 37°C. The next steps including washes, hybridization, ligation and amplification were performed following insert recommendations. Slides were mounted with *in situ* mounting medium with DAPI (Sigma-Aldrich) and visualized by an LSM 700 confocal microscope. The PLA signals per cell were determined in at least 30 cells in maximum projection images using the Spot Detector plugin in Icy Image software (65). Mock-infected cells incubated with both primary antibodies were used as negative controls.

### Western blotting

Cells were treated with lysis buffer (25 mM Tris-HCl, pH 7.6, 150 mM NaCl, 1% Triton X-100, 1% sodium deoxycholate, 0.1% SDS, 5% glycerol) supplemented with protease inhibitor cocktail (P8340; Sigma-Aldrich). Total protein concentration was determinated using Pierce bicinchoninic acid protein assay kit (Thermo Scientific) and 20µg by samples were mixed in Laemmli loading buffer (40% glycerol, 240 mM Tris-HCl, pH 6.8, 8% SDS, 0.04% bromophenol blue, 5% β-mercaptoethanol) at 1X final concentration. Samples were then denatured by 5 minutes in boiling water and loaded in polyacrylamide gels with SDS–10%. After electrophoresis proteins were transferred to the nitrocellulose membrane (0.45 µm; Bio-Rad), and incubated with primary antibodies diluted in 5% skin milk powder in PBS–0.1% Tween 20. After washing, membranes were incubated with secondary antibodies conjugated to HRP (anti-mouse-HRP 115-035-003; 1/20,000; Jackson ImmunoResearch; or anti-rabbit-HRP GTX26821; 1/40,000; Genetex) diluted in 5% skin milk powder in PBS–0.1% Tween 20. HRP was detected using SuperSignal West Femto maximum sensitivity substrate (Thermo Scientific). Digital images were acquired with a Fusion FX Spectra (Vilber) and analyzed with ImageJ software (60). **BioID system.** The protocol used to label NS1 interacting proteins with biotin was as reported by Roux et al (28, 29, 66) small modifications. Transfected mosquito cells were incubated for 18 hours with EMEM 10% SFB supplemented with 50 μM biotin. Quadruplicates of each condition were carried out. After biotinylating time, each well (6-well plate format) was washed twice with PBS. Then, cells were lysed at 25°C in 1 ml lysis buffer (50 mM Tris, pH 7.4, 500 mM NaCl, 0.4% SDS, 5 mM EDTA, 1 mM DTT) supplemented with protease inhibitor cocktail (P8340; Sigma-Aldrich). Lysates were pooled, vortex the 3 times for 1 minute, and incubated with 600 μl of Dynabeads (MyOne Steptavadin C1; Invitrogen) overnight at 4°C with gentle mixing by inversion in a rotor mixer. Beads were collected by low speed centrifugation for 8 min at 25°C (all subsequent steps at 25°C) and washed by low speed centrifugation, twice with 1 ml of wash buffer 1 (2% SDS in dH2O), twice with 1 ml of wash buffer 2 (0.1% deoxycholate, 1% Triton X-100, 500 mM NaCl, 1 mM EDTA, and 50 mM Hepes, pH 7.5), once with 1 ml of wash buffer 3 (250 mM LiCl, 0.5% NP-40, 0.5% deoxycholate, 1 mM EDTA, and 10 mM Tris, pH 8.1) and twice with 1 ml of buffer 4 (50 mM Tris, pH 7.4, and 50 mM NaCl). Finally, samples were washed twice in 50 mM NH_4_HCO_3_ to be analyzed by mass spectrometry.

### Protein identification by mass spectrometry

Sample preparation was carried out following the protocol described by Ren et al. (67) for LC-MS/MS, in combination with an equivalent BioID system, named RNA-protein interaction detection (RaPID). Analysis was conducted in the Stanford University Mass Spectrometry Facility (CA, USA). Streptavidin magnetic beads were re-suspended in 200 μl of 50 mM ammonium bicarbonate supplemented with DTT to a final concentration of 5 mM, incubated on a heat block at 50 °C for 5 min, followed by head over head rocking for 30 min at 25°C. Alkylation was performed by the addition of propionamide to a final concentration of 10 mM and head over head rocking for 30 min at 25°C. Trypsin/LysC (Promega) was added to each sample to a final concentration of 250 ng, and digested overnight at 25°C in the head overhead shaker, followed by the addition of formic acid to 1% final concentration. Peptides were removed and washed with 50 μl of 0.1% formic acid in distilled water. Finally, the acidified peptide pools were purified with C18 STAGE tip 37 (NEST group) microspin columns and dried in a speed-vac. For LCMS/ MS, peptide pools were reconstituted and injected onto a C18 reversed phase analytical column. All MS/MS data was first analyzed in Preview to provide recalibration criteria and then reformatted to MGF before full analysis with Byonic v1.4 (ProteinMetrics). Analyses used .fasta files proteins from *Aedes albopictus*, concatenated with common contaminant human proteins and with DENV proteins. Row data were searched at 10 ppm mass tolerances for precursors, with 0.4 Da fragment mass tolerances assuming up to two missed cleavages and allowing for N-ragged tryptic digestion and were validated at a 1% false discovery rate, using typical reverse-decoy techniques. The resulting assigned proteins were then filtered excluding proteins with less than 2 unique peptides and 6 spectra and exported for further analysis using custom bioinformatic tools.

### Transcriptomics

For mRNA sequencing, C6/36 cells were seeded in 6 well format plates. Assay was performed simultaneously for four conditions: C1) cells mock infected and treated with irrelevant siRNA, C2) cells mock infected and treated with DIDO1 siRNA, C3) cells infected with DENV2 at MOI=3 3 for 18 hours and treated with irrelevant siRNA, C4) cells infected with DENV2 at MOI=3 for 18 hours and treated with DIDO1 siRNA. Three replicates were used for each condition. Gene silencing and infection were performed as previously described. Total RNA was extracted with the RNeasy Mini Kit (QIAGEN., CAT.74104) following insert instructions. Samples were shipped to Theragen (http://www.theragenetex.com/en/) for next generation sequencing (NGS). Sequencing was performed in a Novaseq6000 (Illumina) instrument, paired-end format to obtain up to 70M reads per sample. Total RNA integrity was evaluated using Agilent 2100 BioAnalyzer with RNA Integrity Number (RIN) value greater than 6. Poly-A selection for transcript mRNA enrichment was chosen and Illumina TruSeq mRNA stranded prep kit. Biological replicates were pooled for each condition (C1-C4) to increase reading/mapping quality. For genome assembly Bowtie2 (53), Cufflinks (68) and TopHat2 (54) packages were used. FastQC was used for quality assessment of sequencing data for all samples (https://www.bioinformatics.babraham.ac.uk/projects/fastqc/).

Overall mapping for each condition was low-but-expected for the whole genome (53.71 %, 54.02 %, 53.22 % and 54.01 % mapped reads for C1, C2, C3 and C4, respectively). However, much better mapping was obtained for gene sequences (88.73 %, 89.06 %, 88.90 % and 89.11 % for C1, C2, C3 and C4, respectively). Proportions of properly paired sequences in gene regions were also appropriate enough (86.28 %, 86.03 %, 85.72 % and 85.75 % C1, C2, C3 and C4, respectively). Sequencing depth randomness plots were unimodal and symmetric for all 4 conditions; maximum depth values were between 100 and 130 M in all cases. Aligned reads in the gene regions were all above 60M for the four conditions. Base sequence quality (Phred scores) distributions sharply peaked at Q=36; this is an approximate probability for an incorrect base call of 1 in 3981, well above the Q=30 (1 error in 1000 calls) standard.

Base contents passed-the-barcodes were well-balanced in all four conditions. GC content deviations from the expected theoretical distributions were not statistically significant in any of the four conditions, thus no significant GC-bias is to be expected in transcript counts. No isoform or novel genes were mapped.

Expression coverage for aligned sequence in coding regions was fully obtained in all four conditions. All genes aligned and mapped have transcripts read. A total of 55,180 transcripts, corresponding to 21,441 known genes, were read. Primary transcriptomic analysis was made with Cufflinks (68). The Cufflinks algorithm detects sequence-specific bias and corrects for it in abundance estimation by re-estimating the transcript abundances dividing each multi-mapped read probabilistically based on the initial abundance estimation, the inferred fragment length and fragment bias. Differential expression analysis was performed with the CuffDiff algorithm in Cufflinks and with R using DESeq2 (69), setting p < 0.05. Multidimensional scaling (MDS) with a Euclidean distance was used. Visualization was performed in R using the pheatmap and EnhancedVolcano packages (https://www.bioconductor.org/).

### Statistical analysis

Plotted results, obtained from three independent experiments, are expressed as means ± standard errors. Statistical analyses were carried out using GraphPad Prism, version 6.01, software.

## Acknowledgements

Authors like to thank Dr. Enrique Hernández Lemus for his critical reading of the manuscript. This work was partially funded by CONACYT-Mexico through a doctoral degree scholarship to GC and a basic science research grant (CB-2015-1 number 254461) to JEL. Authors declare no conflict of interest.

## Author contributions statement

GIC and JEL conceived and designed the experiments; GIC, RRR, MV and SD conducted the experiments; all authors analyzed the results; JEL and HBG provided reagents and financial support; GIC wrote the first draft of the manuscript; GIC and JEL wrote the final version of the manuscript; all authors reviewed the manuscript.

---

**Supplemental Figure 1.**
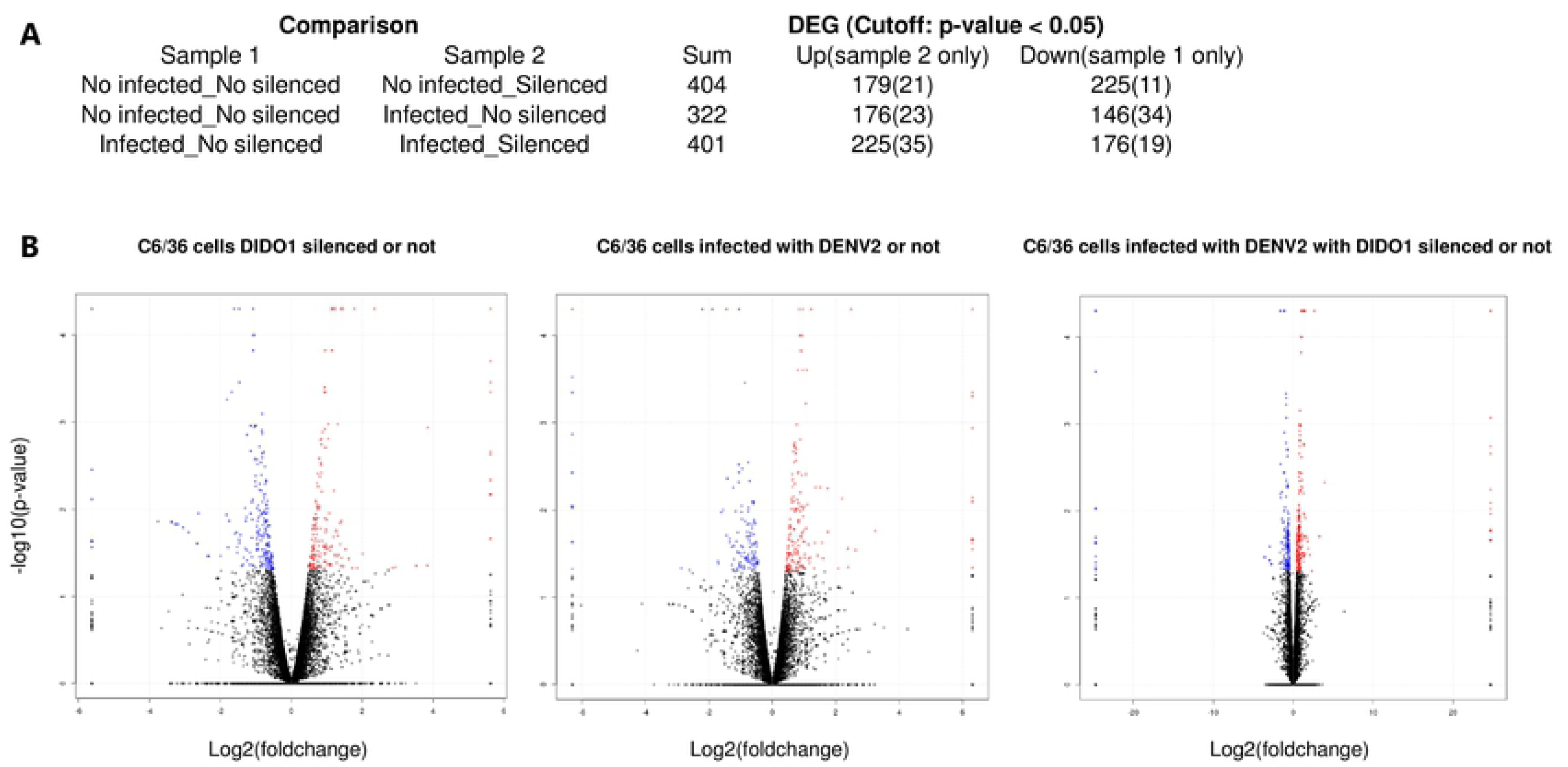
(A) Overall resume of results comparing pairs of conditions as indicated. Combined conditions include infection status with DENV2 (MOI=3) and/or DIDO1 silencing status. DEG= Differential expressed genes; cutoff p-value used was < 0.05. (B) Volcano plots of 3 different conditions as indicated in the top of each plot. Colored dots represent genes with p-value < 0.05, blue = down expressed and red= up expressed.

**Supplemental Figure 2.**
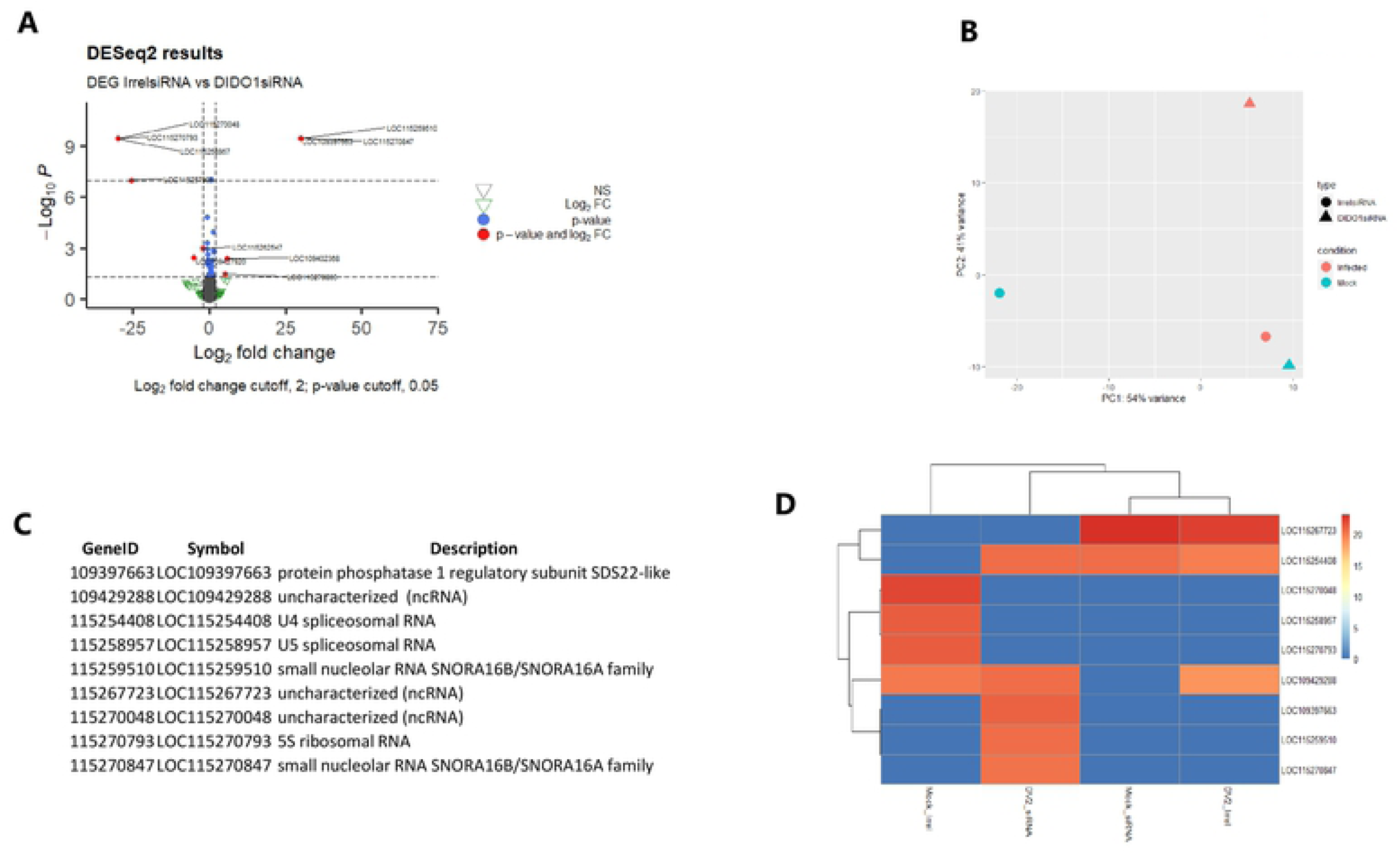
(A) DEG due to Dido1 silencing status, independently of the infection status. Only 10 genes are highlighted due to p-value and log2 fold change cutoffs. (B) PCA plot for conditions comparison and clustering. Interestingly, C6/36 cells infected show better similarity with non-infected cells with DIDO1 silenced. Clustering (C) and heatmap (D) analysis of selected DEG.

## References

1. Holbrook MR. Historical Perspectives on Flavivirus Research https://doi.org/10.3390/v9050097.

2. Zhang Y, Gao W, Li J, Wu W, Jiu Y. 2019. The Role of Host Cytoskeleton in Flavivirus Infection. Virol Sin 34:30–41.

3. Luo D, Xu T, Watson RP, Scherer-Becker D, Sampath A, Jahnke W, Yeong SS, Wang CH, Lim SP, Strongin A, Vasudevan SG, Lescar J. 2008. Insights into RNA unwinding and ATP hydrolysis by the flavivirus NS3 protein. EMBO J 27:3209–3219.

4. Sahili A El, Lescar J. 2017. Dengue virus non-structural protein 5. Viruses. MDPI AG.

5. Li Y, Kim YM, Zou J, Wang QY, Gayen S, Wong YL, Lee LT, Xie X, Huang Q, Lescar J, Shi PY, Kang C. 2015. Secondary structure and membrane topology of dengue virus NS4B N-terminal 125 amino acids. Biochim Biophys Acta - Biomembr 1848:3150–3157.

6. Egloff MP, Decroly E, Malet H, Selisko B, Benarroch D, Ferron F, Canard B. 2007. Structural and Functional Analysis of Methylation and 5′-RNA Sequence Requirements of Short Capped RNAs by the Methyltransferase Domain of Dengue Virus NS5. J Mol Biol 372:723–736.

7. Dong H, Chang DC, Hua MHC, Lim SP, Chionh YH, Hia F, Lee YH, Kukkaro P, Lok SM, Dedon PC, Shi PY. 2012. 2′-O methylation of internal adenosine by flavivirus NS5 methyltransferase. PLoS Pathog 8.

8. Egloff MP, Benarroch D, Selisko B, Romette JL, Canard B. 2002. An RNA cap (nucleoside-2′-O-)-methyltransferase in the flavivirus RNA polymerase NS5: Crystal structure and functional characterization. EMBO J 21:2757–2768.

9. Ackermann M, Padmanabhan R. 2001. De Novo Synthesis of RNA by the Dengue Virus RNA-dependent RNA Polymerase Exhibits Temperature Dependence at the Initiation but Not Elongation Phase. J Biol Chem 276:39926–39937.

10. Zeidler JD, Fernandes-Siqueira LO, Barbosa GM, Da Poian AT. 2017. Non-canonical roles of dengue virus non-structural proteins. Viruses 9.

11. Muller DA, Young PR. 2013. The flavivirus NS1 protein: Molecular and structural biology, immunology, role inpathogenesis and application asadiagnostic biomarker. Antiviral Res. Elsevier B.V.

12. Chen H, Lai Y, Yeh T. 2018. Dengue virus non-structural protein 1 : a pathogenic factor, therapeutic target , and vaccine candidate 1–11.

13. Mackenzie JM, Jones MK, Young PR. 1996. Immunolocalization of the Dengue virus nonstructural glycoprotein NS1 suggests a role in viral RNA replication. Virology 220:232–240.

14. Westaway EG, Mackenzie JM, Kenney MT, Jones MK, Khromykh AA. 1997. Ultrastructure of Kunjin virus-infected cells: colocalization of NS1 and NS3 with double-stranded RNA, and of NS2B with NS3, in virus-induced membrane structures. J Virol 71:6650–6661.

15. MacKenzie JM, Khromykh AA, Jones MK, Westaway EG. 1998. Subcellular localization and some biochemical properties of the flavivirus Kunjin nonstructural proteins NS2A and NS4A. Virology 245:203–215.

16. Edeling MA, Diamond MS, Fremont DH. 2014. Structural basis of flavivirus NS1 assembly and antibody recognition. Proc Natl Acad Sci U S A 111:4285–4290.

17. Gutsche I, Coulibaly F, Voss JE, Salmon J, D’Alayer J, Ermonval M, Larquet E, Charneau P, Krey T, Mégret F, Guittet E, Rey FA, Flamand M. 2011. Secreted dengue virus nonstructural protein NS1 is an atypical barrel-shaped high-density lipoprotein. Proc Natl Acad Sci U S A 108:8003–8008.

18. Muller DA, Young PR. 2013. The flavivirus NS1 protein: Molecular and structural biology, immunology, role inpathogenesis and application asadiagnostic biomarker. Antiviral Res 98:192–208.

19. Liu Y, Liu J, Du S, Shan C, Nie K, Zhang R, Li XF, Zhang R, Wang T, Qin CF, Wang P, Shi PY, Cheng G. 2017. Evolutionary enhancement of Zika virus infectivity in Aedes aegypti mosquitoes. Nature 545:482–486.

20. Hafirassou ML, Meertens L, Umaña-Diaz C, Labeau A, Dejarnac O, Bonnet-Madin L, Kümmerer BM, Delaugerre C, Roingeard P, Vidalain P-OO, Amara A. 2017. A Global Interactome Map of the Dengue Virus NS1 Identifies Virus Restriction and Dependency Host Factors. Cell Rep 21:3900–3913.

21. Rastogi M, Sharma N, Singh SK. 2016. Flavivirus NS1: a multifaceted enigmatic viral protein. Virol J 13:131.

22. Glasner DR, Puerta-Guardo H, Beatty PR, Harris E. 2018. The Good, the Bad, and the Shocking: The Multiple Roles of Dengue Virus Nonstructural Protein 1 in Protection and Pathogenesis. Annu Rev Virol 5:annurev-virology-101416-041848.

23. Alcalá AC, Palomares LA, Ludert JE. 2018. Secretion of Nonstructural Protein 1 of Dengue Virus from Infected Mosquito Cells: Facts and Speculations. J Virol 92.

24. Alcalá AC, Medina F, González-Robles A, Salazar-Villatoro L, Fragoso-Soriano RJ, Vásquez C, Cervantes-Salazar M, del Angel RM, Ludert JE. 2016. The dengue virus non-structural protein 1 (NS1) is secreted efficiently from infected mosquito cells. Virology 488:278–287.

25. Reyes-Ruiz JM, Osuna-Ramos JF, Cervantes-Salazar M, Lagunes Guillen AE, Chávez-Munguía B, Salas-Benito JS, Del Ángel RM. 2018. Strand-like structures and the nonstructural proteins 5, 3 and 1 are present in the nucleus of mosquito cells infected with dengue virus. Virology 515:74–80.

26. Alcalá AC, Hernández-Bravo R, Medina F, Coll DS, Zambrano JL, Del Angel RM, Ludert JE. 2017. The dengue virus non-structural protein 1 (NS1) is secreted from infected mosquito cells via a non-classical caveolin-1-dependent pathway. J Gen Virol 98:2088–2099.

27. Rosales Ramirez R, Ludert JE. 2019. The Dengue Virus Nonstructural Protein 1 (NS1) Is Secreted from Mosquito Cells in Association with the Intracellular Cholesterol Transporter Chaperone Caveolin Complex. J Virol 93.

28. Roux KJ, Kim DI, Raida M, Burke B. 2012. A promiscuous biotin ligase fusion protein identifies proximal and interacting proteins in mammalian cells. J Cell Biol 196:801–810.

29. Roux KJ, Kim DI, Burke B, May DG. 2018. BioID: A Screen for Protein-Protein Interactions. Curr Protoc protein Sci 91:19.23.1–19.23.15.

30. González M, Martín-Ruíz I, Jiménez S, Pirone L, Barrio R, Sutherland JD. 2011. Generation of stable Drosophila cell lines using multicistronic vectors. Sci Rep 1:75.

31. Avirutnan P, Fuchs A, Hauhart RE, Somnuke P, Youn S, Diamond MS, Atkinson JP. 2010. Antagonism of the complement component C4 by flavivirus nonstructural protein NS1. J Exp Med 207:793–806.

32. Shah PS, Link N, Jang GM, Sharp PP, Zhu T, Swaney DL, Johnson JR, Von Dollen J, Ramage HR, Satkamp L, Newton B, Hüttenhain R, Petit MJ, Baum T, Everitt A, Laufman O, Tassetto M, Shales M, Stevenson E, Iglesias GN, Shokat L, Tripathi S, Balasubramaniam V, Webb LG, Aguirre S, Willsey AJ, Garcia-Sastre A, Pollard KS, Cherry S, Gamarnik A V., Marazzi I, Taunton J, Fernandez-Sesma A, Bellen HJ, Andino R, Krogan NJ. 2018. Comparative Flavivirus-Host Protein Interaction Mapping Reveals Mechanisms of Dengue and Zika Virus Pathogenesis. Cell 175:1931–1945.

33. Noisakran S, Sengsai S, Thongboonkerd V, Kanlaya R, Sinchaikul S, Chen S-TT, Puttikhunt C, Kasinrerk W, Limjindaporn T, Wongwiwat W, Malasit P, Yenchitsomanus P thai. 2008. Identification of human hnRNP C1/C2 as a dengue virus NS1-interacting protein. Biochem Biophys Res Commun 372:67–72.

34. Avirutnan P, Hauhart RE, Somnuke P, Blom AM, Diamond MS, Atkinson JP. 2011. Binding of Flavivirus Nonstructural Protein NS1 to C4b Binding Protein Modulates Complement Activation. J Immunol 187:424–433.

35. Allonso D, Andrade IS, Conde JN, Coelho DR, Rocha DCP, da Silva ML, Ventura GT, Silva EM, Mohana-Borges R. 2015. Dengue Virus NS1 Protein Modulates Cellular Energy Metabolism by Increasing Glyceraldehyde-3-Phosphate Dehydrogenase Activity. J Virol 89:11871–83.

36. Rabelo K, Trugilho MROO, Costa SM, Pereira BASS, Moreira OC, Ferreira ATSS, Carvalho PC, Perales J, Alves AMBB. 2017. The effect of the dengue non-structural 1 protein expression over the HepG2 cell proteins in a proteomic approach. J Proteomics 152:339–354.

37. Dechtawewat T, Paemanee A, Roytrakul S, Songprakhon P, Limjindaporn T, Yenchitsomanus P thai, Saitornuang S, Puttikhunt C, Kasinrerk W, Malasit P, Noisakran S. 2016. Mass spectrometric analysis of host cell proteins interacting with dengue virus nonstructural protein 1 in dengue virus-infected HepG2 cells. Biochim Biophys Acta - Proteins Proteomics https://doi.org/10.1016/j.bbapap.2016.04.008.

38. Songprakhon P, Limjindaporn T, Perng GC, Puttikhunt C, Thaingtamtanha T, Dechtawewat T, Saitornuang S, Uthaipibull C, Thongsima S, Yenchitsomanus PT, Malasit P, Noisakran S. 2018. Human glucose-regulated protein 78 modulates intracellular production and secretion of nonstructural protein 1 of dengue virus. J Gen Virol 99:1391–1406.

39. Doolittle JM, Gomez SM. 2011. Mapping protein interactions between dengue virus and its human and insect hosts. PLoS Negl Trop Dis 5.

40. Khadka S, Vangeloff AD, Zhang C, Siddavatam P, Heaton NS, Wang L, Sengupta R, Sahasrabudhe S, Randall G, Gribskov M, Kuhn RJ, Perera R, Lacount DJ. 2011. A Physical Interaction Network of Dengue Virus and Human Proteins https://doi.org/10.1074/mcp.M111.012187.

41. Cervantes-Salazar M, Angel-Ambrocio AH, Soto-Acosta R, Bautista-Carbajal P, Hurtado-Monzon AM, Alcaraz-Estrada SL, Ludert JE, Del Angel RM. 2015. Dengue virus NS1 protein interacts with the ribosomal protein RPL18: This interaction is required for viral translation and replication in Huh-7 cells. Virology https://doi.org/10.1016/j.virol.2015.05.017.

42. Warde-Farley D, Donaldson SL, Comes O, Zuberi K, Badrawi R, Chao P, Franz M, Grouios C, Kazi F, Lopes CT, Maitland A, Mostafavi S, Montojo J, Shao Q, Wright G, Bader GD, Morris Q. The GeneMANIA prediction server: biological network integration for gene prioritization and predicting gene function https://doi.org/10.1093/nar/gkq537.

43. von Mering C, Jensen LJ, Snel B, Hooper SD, Krupp M, Foglierini M, Jouffre N, Huynen MA, Bork P. 2005. STRING: Known and predicted protein-protein associations, integrated and transferred across organisms. Nucleic Acids Res 33:D433.

44. Shannon P, Markiel A, Ozier O, Baliga NS, Wang JT, Ramage D, Amin N, Schwikowski B, Ideker T. 2003. Cytoscape: A software Environment for integrated models of biomolecular interaction networks. Genome Res 13:2498–2504.

45. Garcia-Domingo D, Ramirez D, Gonzalez de Buitrago G, Martinez-A C. 2003. Death Inducer-Obliterator 1 Triggers Apoptosis after Nuclear Translocation and Caspase Upregulation. Mol Cell Biol 23:3216–3225.

46. Fütterer A, de Celis J, Navajas R, Almonacid L, Gutiérrez J, Talavera-Gutiérrez A, Pacios-Bras C, Bernascone I, Martin-Belmonte F, Martinéz-A C. 2017. DIDO as a Switchboard that Regulates Self-Renewal and Differentiation in Embryonic Stem Cells. Stem Cell Reports 8:1062–1075.

47. Liu Y, Kim H, Liang J, Lu W, Ouyang B, Liu D, Songyang Z. 2014. The death-inducer obliterator 1 (Dido1) gene regulates embryonic stem cell self-renewal. J Biol Chem 289:4778–4786.

48. Mora Gallardo C, Sánchez de Diego A, Gutiérrez Hernández J, Talavera-Gutiérrez A, Fischer T, Martínez-A C, van Wely KHM. 2019. Dido3-dependent SFPQ recruitment maintains efficiency in mammalian alternative splicing. Nucleic Acids Res 47:5381–5394.

49. Bindea G, Mlecnik B, Hackl H, Charoentong P, Tosolini M, Kirilovsky A, Fridman WH, Pagès F, Trajanoski Z, Galon J. 2009. ClueGO: A Cytoscape plug-in to decipher functionally grouped gene ontology and pathway annotation networks. Bioinformatics 25:1091–1093.

50. Xin Q-L, Deng C-L, Chen X, Wang J, Wang S-B, Wang W, Deng F, Zhang B, Xiao G, Zhang L-K, Xin Q-L C, C-l D, S-b W, Michael Diamond ES. 2017. Quantitative Proteomic Analysis of Mosquito C6/36 Cells Reveals Host Proteins Involved in Zika Virus Infection Downloaded from. jvi.asm.org 1 J Virol 91:554–571.

51. Li M-J, Lan C-J, Gao H-T, Xing D, Gu Z-Y, Su D, Zhao T-Y, Yang H-Y, Li C-X. Transcriptome analysis of Aedes aegypti Aag2 cells in response to dengue virus-2 infection https://doi.org/10.1186/s13071-020-04294-w.

52. García-Domingo D, Ramírez D, González de Buitrago G, Martínez-A C. 2003. Death Inducer-Obliterator 1 Triggers Apoptosis after Nuclear Translocation and Caspase Upregulation. Mol Cell Biol 23:3216–3225.

53. Tham H-W, Balasubramaniam VR, Chew M-F, Ahmad H, Hassan SS. 2015. Protein-protein interactions between A. aegypti midgut and dengue virus 2: two-hybrid screens using the midgut cDNA library. J Infect Dev Ctries 9:1338–49.

54. Noisakran S, Sengsai S, Thongboonkerd V, Kanlaya R, Sinchaikul S, Chen ST, Puttikhunt C, Kasinrerk W, Limjindaporn T, Wongwiwat W, Malasit P, Yenchitsomanus P thai. 2008. Identification of human hnRNP C1/C2 as a dengue virus NS1-interacting protein. Biochem Biophys Res Commun 372:67–72.

55. Guo X, Xu Y, Bian G, Pike AD, Xie Y, Xi Z. 2010. Response of the mosquito protein interaction network to dengue infection. BMC Genomics 11:1–15.

56. Doolittle JM, Gomez SM, Fc D V, Angleró-rodríguez YI, Macleod HJ, Kang S, Carlson JS, Jupatanakul N, Dimopoulos G, Huang YJS, Higgs S, Horne KME, Vanlandingham DL, Doolittle JM, Gomez SM, Guo X, Xu Y, Bian G, Pike AD, Xie Y, Xi Z. 2011. Mapping protein interactions between dengue virus and its human and insect hosts. PLoS Negl Trop Dis 5:4703–4730.

57. De Maio FA, Risso G, Iglesias NG, Shah P, Pozzi B, Gebhard LG, Mammi P, Mancini E, Yanovsky MJ, Andino R, Krogan N, Srebrow A, Gamarnik A V. 2016. The Dengue Virus NS5 Protein Intrudes in the Cellular Spliceosome and Modulates Splicing. PLoS Pathog 12:1–29.

58. Braig S, Bosserhoff AK. 2013. Death inducer-obliterator 1 (Dido1) is a BMP target gene and promotes BMP-induced melanoma progression. Oncogene 32:837–848.

59. Johnson ML, Nagengast AA, Salz HK. 2010. PPS, a Large Multidomain Protein, Functions with Sex-Lethal to Regulate Alternative Splicing in Drosophila. PLoS Genet 6:1000872.

60. Rueden CT, Schindelin J, Hiner MC, DeZonia BE, Walter AE, Arena ET, Eliceiri KW. 2017. ImageJ2: ImageJ for the next generation of scientific image data. BMC Bioinformatics 18:529.

61. Dzaki N, Azzam G. 2018. Assessment of aedes albopictus reference genes for quantitative PCR at different stages of development. PLoS One 13:1–19.

62. Chien LJ, Liao TL, Shu PY, Huang JH, Gubler DJ, Chang GJJ. 2006. Development of real-time reverse transcriptase PCR assays to detect and serotype dengue viruses. J Clin Microbiol 44:1295–1304.

63. Lanciotti RS, Kosoy OL, Laven JJ, Velez JO, Lambert AJ, Johnson AJ, Stanfield SM, Duffy MR. 2008. Genetic and serologic properties of Zika virus associated with an epidemic, Yap State, Micronesia, 2007. Emerg Infect Dis 14:1232–1239.

64. Biering SB, Akey DL, Wong MP, Clay Brown W, Lo NTN, Puerta-Guardo H, de Sousa FTG, Wang C, Konwerski JR, Espinosa DA, Bockhaus NJ, Glasner DR, Li J, Blanc SF, Juan EY, Elledge SJ, Mina MJ, Robert Beatty P, Smith JL, Harris E. 2021. Structural basis for antibody inhibition of flavivirus NS1–triggered endothelial dysfunction. Science (80-) 371:194–200.

65. De Chaumont F, Dallongeville S, Chenouard N, Hervé N, Pop S, Provoost T, Meas-Yedid V, Pankajakshan P, Lecomte T, Le Montagner Y, Lagache T, Dufour A, Olivo-Marin JC. 2012. Icy: An open bioimage informatics platform for extended reproducible research. Nat Methods. Nature Publishing Group.

66. Kim DI, Jensen SC, Noble KA, KC B, Roux KH, Motamedchaboki K, Roux KJ. 2016. An improved smaller biotin ligase for BioID proximity labeling. Mol Biol Cell 27:1188–1196.

67. Ren L, Ding S, Song Y, Li B, Ramanathan M, Co J, Amieva MR, Khavari PA, Greenberg HB. 2019. Profiling of rotavirus 3UTR-binding proteins reveals the ATP synthase subunit ATP5B as a host factor that supports late-stage virus replication. J Biol Chem 294:5993–6006.

68. Trapnell C, Williams BA, Pertea G, Mortazavi A, Kwan G, Van Baren MJ, Salzberg SL, Wold BJ, Pachter L. 2010. Transcript assembly and quantification by RNA-Seq reveals unannotated transcripts and isoform switching during cell differentiation. Nat Biotechnol 28:511–515.

69. Love MI, Huber W, Anders S. 2014. Moderated estimation of fold change and dispersion for RNA-seq data with DESeq2. Genome Biol 15:550.

